# Twisting Urea- to Imide-Based Mass Spectrometry-Cleavable Cross-Linkers Enables Affinity Tagging

**DOI:** 10.1101/2024.03.29.587196

**Authors:** Alessio Di Ianni, Christian H. Ihling, Tomáš Vranka, Václav Matoušek, Andrea Sinz, Claudio Iacobucci

## Abstract

Disuccinimidyl dibutyric urea (DSBU) is a mass spectrometry (MS)-cleavable cross-linker that has multiple applications in structural biology, ranging from isolated protein complexes to comprehensive system-wide interactomics. DSBU facilitates a rapid and reliable identification of cross-links through the dissociation of its urea group in the gas-phase. In this study, we further advance the structural capabilities of DSBU by twisting the urea group into an imide, thus introducing a novel class of cross-linkers. This modification preserves the MS-cleavability of the amide bond, granted by the two acyl groups of the imide function. The central nitrogen atom enables the introduction of affinity purification tags. Here, we introduce disuccinimidyl disuccinic imide (DSSI) as prototype of this class of cross-linkers. It features a phosphonate handle for immobilized metal ion affinity chromatography (IMAC) enrichment. We detail DSSI synthesis and describe its behavior in solution and in the gas-phase while cross-linking isolated proteins and human cell lysates. DSSI and DSBU cross-links are compared at the same enrichment depths to bridge these two cross-linker classes. We validate DSSI cross-links by mapping them in high-resolution structures of large protein assemblies. The cross-links observed yield insights into the morphology of intrinsically disordered proteins (IDPs) and their complexes. The DSSI linker might spearhead a novel class of MS-cleavable and enrichable cross-linkers.

## Introduction

Over the past two decades, cross-linking mass spectrometry (XL-MS) has emerged as a powerful technique for exploring the 3D-structures of proteins and protein complexes in their native environment [1, 2]. Chemical cross-linkers covalently bridge two amino acids in spatial proximity and serve as molecular rulers to unveil the morphology and plasticity of proteins. The major bottleneck of XL-MS remains the analysis of the complex enzymatic proteolysates obtained from the cross-linked sample. Cross-links are formed sub-stoichiometrically and are dispersed in a several orders of magnitude higher concentrated background of tryptic peptides. The direct application of reverse-phase (RP) liquid chromatography coupled to tandem mass spectrometry (LC-MS/MS) does not provide a comprehensive identification of cross-links, especially in system-wide XL-MS experiments [3]. Adding further chromatographic dimensions enhances the depth of the analysis. Size exclusion (SEC) [4], strong cation-exchange (SCX) [5], hSAX [6], and high-pH RP [7] chromatography leverage the increased average size and charge state of cross-links over linear peptides to pre-fractionate proteolyzed cross-linked samples [8]. Also, several cross-linkers incorporate molecular tags like biotin, azide [9], alkyne [9, 10], or phosphonate [11], enabling affinity purification of cross-links. The combination of affinity purification followed by SEC or SCX allows reducing the background of linear peptides in the sample [12-14]. Among the affinity handles, the phosphonate has recently emerged as one of the more promising options in system-wide XL-MS [12]. Phosphonate-containing cross-links can be enriched by IMAC or on titanium dioxide (TiO_2_) beads. This approach is derived from conventional phosphoproteomics and, unlike phosphopeptides, phosphonates do not suffer from chemical or enzymatic instability. The complexity of cross-linked samples also affects downstream MS data analysis. Here, MS-cleavable cross-linkers like DSBU [15, 16], DSSO [17], PIR [18], and others represent a valid solution. By dissociating in the gas-phase, they reveal the individual masses of the cross-linked peptides, reducing the software search space, and improve the sequencing of both peptides [19-21]. In this work, we twisted the urea group of urea-based cross-linkers to an imide (Figure 1). This preserves the MS-cleavability of imide-based reagents while the central nitrogen atom can be functionalized with affinity tags for cross-link enrichment. We introduce DSSI, a novel trifunctional MS-cleavable reagent containing a phosphonate handle for future IMAC enrichment of cross-links. We describe DSSI synthesis and studied in detail its behavior in the gas-phase. We probed DSSI’s reactivity in solution by cross-linking the intrinsically disordered protein α-synuclein (α-syn), and a HEK293T cell lysate. We compared DSSI with DSBU at the same cross-link enrichment depth finding complementary structural information and protein-protein interactions. DSSI cross-links underwent structure-based validation and yielded structural information on intrinsically disordered proteins (IDPs).

**Figure 1:**
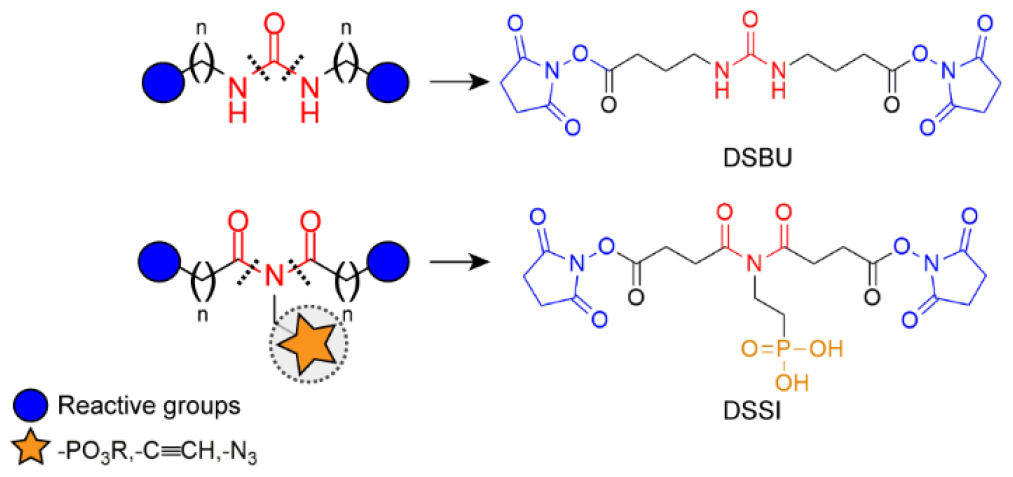
MS-cleavable urea (DSBU) and imide (DSSI) cross-linkers.

## Results and discussions

### Design and synthesis of the DSSI chemical cross-linker

Classical urea-based cross-linkers, such as DSBU, are unsuitable for accommodating an affinity tag without disrupting the molecule’s symmetry. To overcome this limitation, we twisted the central urea group into an imide. This enables the design of cross-linkers featuring a central trivalent nitrogen atom, ideal for attaching affinity handles. At the same time imide-based cross-linkers preserve the MS-cleavability and the spacer length modularity of their urea-based ancestors. DSSI, the first imide-based cross-linker, possesses the same C4 arms of DSBU and a central ethylene phosphonic acid group as affinity tag. DSSI was synthesized using 2-amino ethanol **1** as starting material (Figure 2). **1** was Boc-protected yielding **2** and used to prepare the cyclic ethylene sulfamidate **3**. Dibenzylphosphite sodium salt **4** was subsequently reacted with **3** to obtain the tert-butyl (2-(bis(benzyloxy)phosphoryl)ethyl)carbamate **5**. An acid catalyzed deprotection resulted in the TFA salt **6**. The latter was acylated and coupled to an imidazolide-activated tert-butyl succinic acid yielding the amide **7**. After this step, the NHS ester of tert-butyl succinate was used to obtain the di-tert-butyl imide derivative **8**. TFA deprotection of the tert-butyl groups yielded the diacid **9** which was esterified with N-hydroxysuccinimide (NHS) in the presence of EDC. The resulting dibenzylphosphonic ester **10** was hydrogenated yielding the phosphonic acid **11**, referred to as DSSI here. Synthesis details are provided in the Supporting Information (Figures S1 and S2).

**Figure 2:**
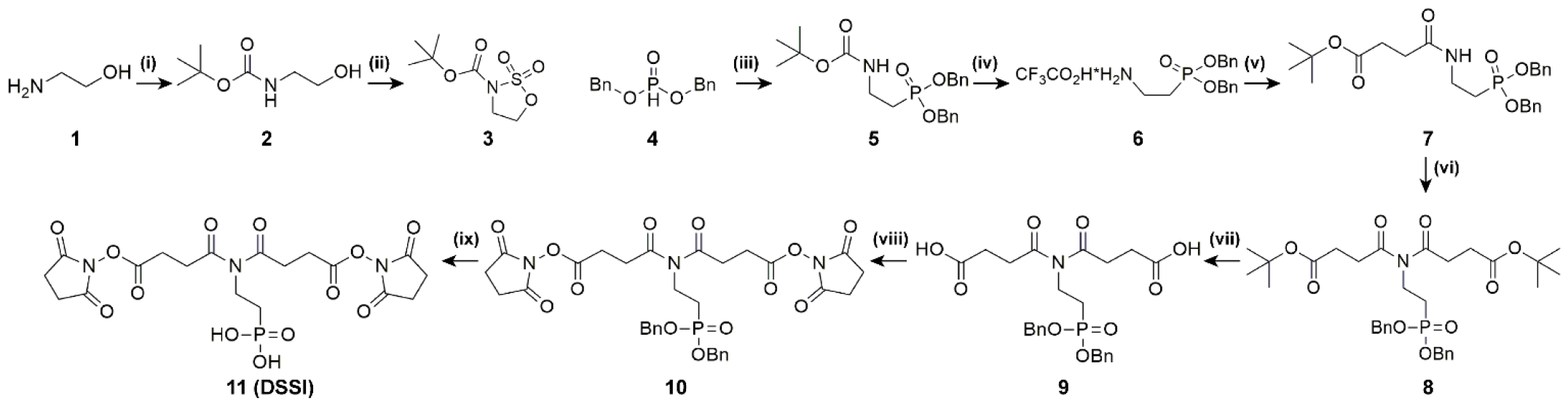
Synthesis of the DSSI cross-linker. i) TEA, DCM, r.t.; ii) Pyridine, CH_3_CN; NaIO_4_, RuCl_3_; iii) **3**, NaH, THF; iv) TFA, DCM; v) butyl succinic acid, CDI, DCM; vi) LiHDMS, -78°C, THF; Succinimidyl mono-t-butyl succinate, THF; vii) TFA, DCM, r.t.; viii) NHS-OH, EDC·HCl, DCM, r.t.; ix) Pd/C, H_2_, THF.

### Characterization of the dissociation behavior of DSSI upon collisional activation

DSBU cross-linked products are cleaved upon collisional activation during MS/MS experiments, resulting in the cleavage of the N-CO bonds of the urea group. The dissociation generates an amine and an isocyanate fragment per peptide visible in fragment ion spectra as two doublets with a characteristic mass difference of ∼26 u [15, 16]. Gas-phase dissociation of linear imides has not been systematically investigated previously. Nelson and McCloskey report that, upon collisional activation, uracil and its derivatives [22] undergo ring-opening via N-CO bond cleavage yielding an amide and a protonated acylium ion. We hypothesized DSSI to follow a similar fragmentation resulting in an MS-cleavable cross-linker. We applied DSSI to cross-link Test Peptide 1 (Ac-TRTESTDIKRASSREADYLINKER, Creative Molecules Inc.) and investigated its dissociation pattern upon collisional activation (Figure S3). Similarly to cyclic imides, DSSI dissociates into ethylphosphonate amides and acylium ions (Figure 3, Figure S4A). Furthermore, one hydroxy group of the phosphonic acid rearranges intramolecularly yielding the dehydrated form of the ethylphosphonate amide fragment and a carboxylic acid fragment for each peptide (Figures 3 and S4B). Thus, DSSI cross-links generate two diagnostic doublets of signals in MS/MS experiments, spaced by ∼89 and ∼125 u (Figure 3). MS/MS experiments of DSSI cross-links revealed that the optimal collision energy for linker and backbone fragmentation ranges between 25 and 35 percent normalized collision energy (NCE), similar to that of DSBU (data not shown). The comparable stabilities of the imide group of DSSI and of the amide bonds in the peptide backbone enable XL-MS analysis at the MS/MS level. This has been shown to increase scan rate and sensitivity compared to multistage MS (MS^n^) methods [23].

**Figure 3:**
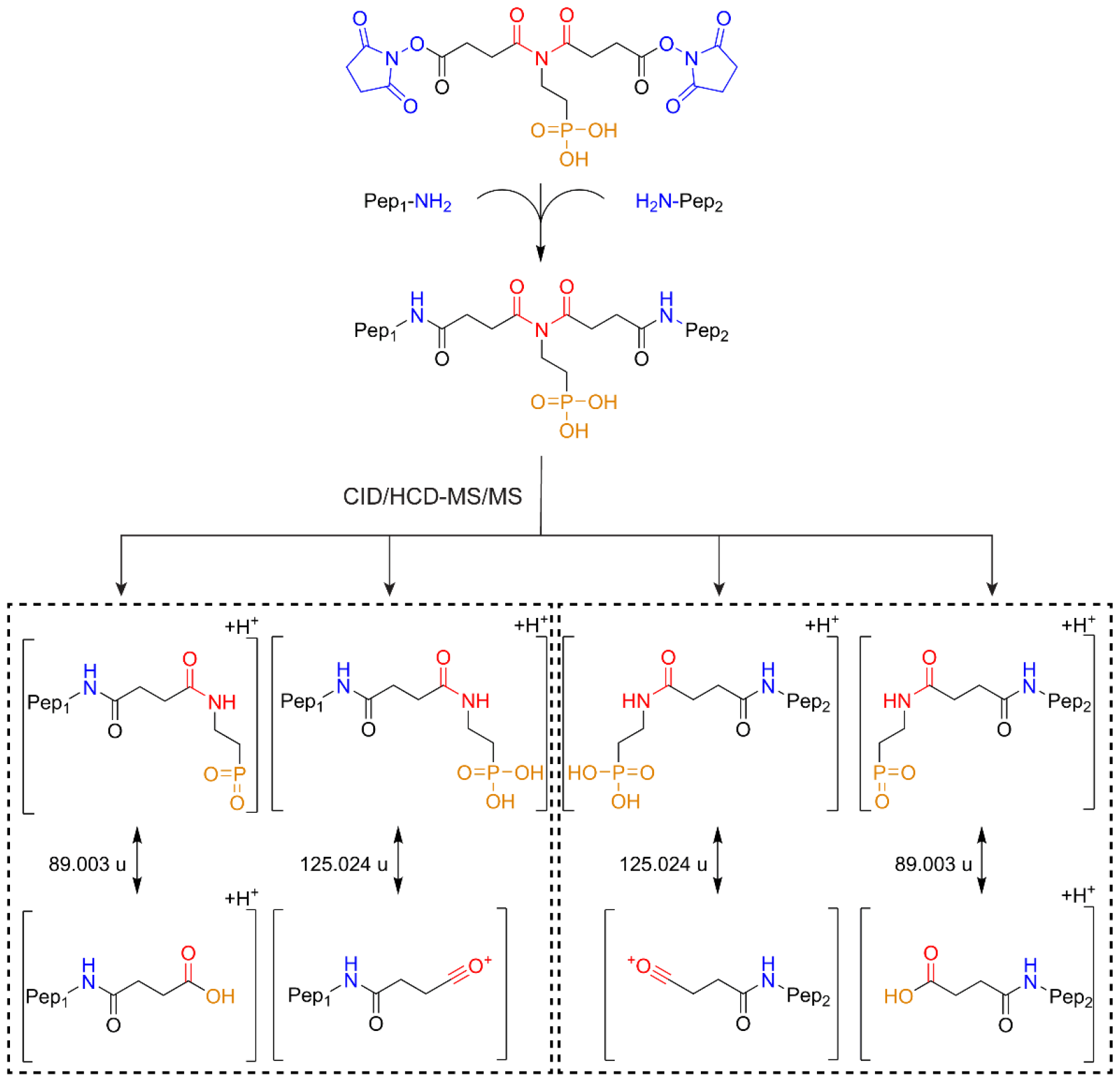
Dissociation behavior of DSSI cross-links upon collisional activation. After the cross-linking reaction, two peptides in spatial proximity are covalently bridged by DSSI. Upon collisional activation, the central imide group dissociates, generating two sets of fragment ions for each cross-linked peptide. These doublets of signals in the fragment ion spectra have a mass difference of ∼125 and ∼89 u.

Additionally, DSSI generates characteristic reporter ions in the low *m/z* range, further enhancing confidence in cross-link identification (Figure S5).

### Application of DSSI for studying single proteins

The thorough characterization of DSSI gas-phase behavior enables the automated cross-link identification in complex samples by using the MeroX software [24]. We selected α-synuclein (α-syn) for assessing the reactivity of DSSI at physiological pH and to establish the downstream sample preparation workflow. α-syn is a small (14.5 kDa) IDP relevant in Parkinson’s disease [25]. It consists of 140 amino acids and three domains, namely the N-terminal lipid-binding domain, the central non-amyloid component (NAC) domain and C-terminal acidic domain. Upon α-syn cross-linking with DSSI, we applied in solution proteolysis for two hours with two consecutive additions of trypsin (fast ISD) [26], and the resulting peptide mixture was analyzed by nanoLC-MS/MS (see Supporting Information for the experimental details). A MeroX search identified 77 unique residue pairs with ∼84% overlap across three replicates (Figure 4B).These were located within the N-terminal domain as well as between the N- and C-terminal regions of α-syn as has been observed with DSBU in our previous study (Figure 4A) [27]. This suggests a comparable reactivity of the two cross-linkers. The high number of DSSI cross-links was achieved despite the application of fast ISD, while DSBU samples underwent overnight digestion (overnight ISD). We adopted the fast ISD for further DSSI cross-linking experiments.

**Figure 4:**
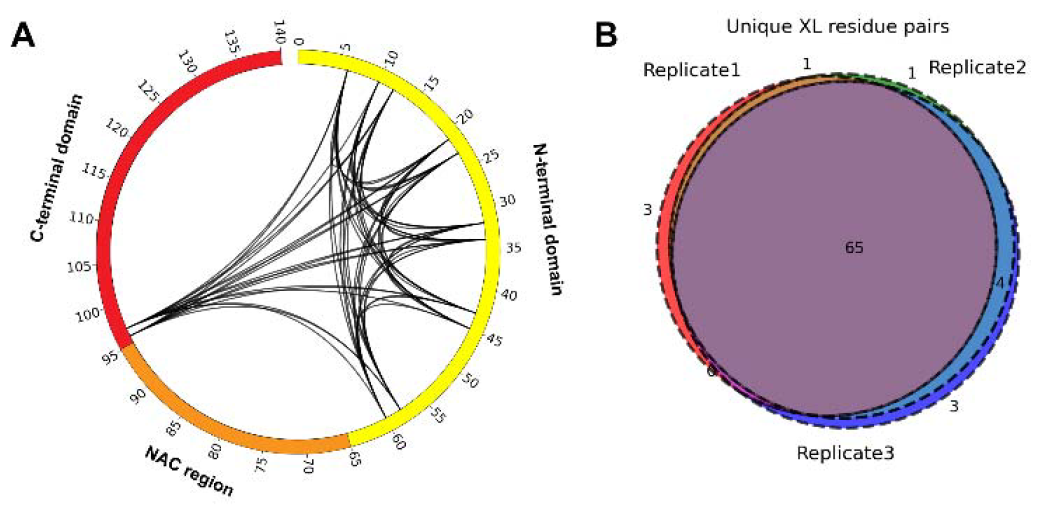
Application of DSSI for studying the intrinsically disordered α-synuclein. A) Circos plot showing identified DSSI cross-links for α-synuclein. B) Reproducibility of cross-linked residue pairs (Venn diagram).

### Application of DSSI cross-linking to cell lysates

Before applying our protocol to more complex samples, we further evaluated fast ISD for HEK293T cell lysates and compared it to a standard overnight ISD. Both differently proteolyzed samples were separated by reverse phase nano-HPLC and analyzed on a timsTOF Pro mass spectrometer (Figure S6). The fast ISD workflow enabled the identification of 5,363 protein groups with 81% reproducibility over three replicates. This corresponds to only 6% fewer protein groups than those identified upon overnight ISD (5,696 protein groups, corresponding to ∼80% reproducibility across three replicates) (Figure S6). The number of identified peptides was comparable between the two workflows (Figure S6 B-C). Therefore, we decided to apply fast ISD to DSSI cross-linked samples to maximize the analysis throughput. We anticipate that reducing the time required for proteolysis will especially benefit the analysis throughput when including the affinity purification of imide-based cross-links. The latter will be addressed in a separate work. Here, our aim was to provide a comprehensive description of DSSI chemistry, comparing it with the well-characterized DSBU. For this reason, DSSI and DSBU cross-links were compared at the same enrichment depth, avoiding the IMAC/TiO_2_ purification step of DSSI cross-links. Cross-linking of HEK293T cell lysates was performed at a total protein concentration of ∼1 g/L and 2 mM DSSI (see Supporting Information for the experimental details). The resulting cross-links were compared to those observed when using DSBU and overnight ISD. Proteolyzed samples were fractionated by SEC to enrich cross-linked peptides (Figure S7A). Selected SEC fractions of each replicate were collected and individually subjected to LC-MS/MS analysis. MeroX analysis allowed the identification of 5,602 unique DSSI cross-links related to 1,241 proteins at 1% false discovery rate (FDR) for cross-link spectrum matches (XSM). Cross-links were non-uniformly distributed over the SEC fractions analyzed, ranging from 147 to 1,440 cross-links identified (Figure S7B). DSSI cross-links provide structural information on 582 protein-protein interactions (PPIs) in HEK293T cells. We compared DSSI cross-linked proteins to those targeted by DSBU for the HEK293T cell line. For this, proteins cross-linked by DSSI and DSBU were analyzed by the clusterProfiler R package [28] and classified based on Gene Ontology (GO) [29]. DSSI- and DBSU-cross-linked proteins were associated with the same cellular compartments of HEK293T cells, consistent with similar properties of the two cross-linkers (Figure S8A-B). In the case of molecular functions, proteins involved in chromatin, tubulin and mRNA/rRNA binding were preferentially cross-linked by DSSI while structural constituents of cytoskeleton, proteins involved in ribonucleoprotein complex binding, NAD binding and RNA helicases were preferentially cross-linked by DSBU (Figure S8C-D). By combining DSSI with DSBU, the cross-linked proteome increases by 18% compared to DSBU alone (Figure S9), enhancing the depth of structural data and the coverage of the PPI network in cells.

DSSI has a similar spacer length as DSBU and is expected to covalently bridge residues with a maximum Cα-Cα distance of 35 Å [30]. We performed a structural validation of DSSI cross-links by mapping them into existing 3D models of three large protein complexes, namely the chaperonin containing TCP-1 (CCT) complex, the 26S proteasome and the 80S ribosome (Figure 5A). The median distances observed ranged between 12 and 22 Å for both cross-linkers. 99% of DSSI cross-links were compatible with known high-resolution structures of the three protein assemblies (Figure 4B).

**Figure 5:**
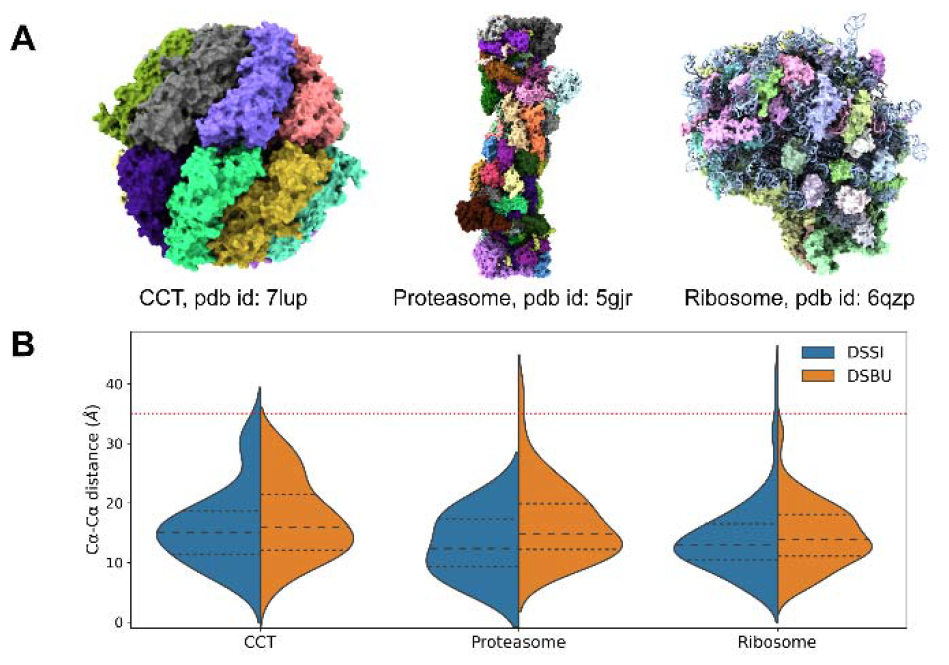
Structural XL-MS validation of DSSI cross-links. A) Selected protein complexes: Chaperonin Containing TCP-1 (CCT), ∼950 kDa; 26S Proteasome, ∼2.5 MDa; 80S Ribosome, ∼4.5 MDa. B) Violin plots comparing the Cα-Cα distance distributions of DSSI (blue) and DSBU (orange) cross-links. Dashed and dotted lines represent the median and the interquartile ranges, respectively.

### System-wide cross-linking of intrinsically disordered proteins

To further evaluate the structural information recapitulated by our cross-linking data, we focused on IDPs. Up to half of all human proteins contain structurally disordered regions (IDRs), which present challenges for visualization using high-resolution methods. [31] The characterization of their highly dynamic structural ensembles requires integrative approaches, including XL-MS experiments. The key role of XL-MS in capturing the plasticity of IDPs is testified by our experiments on HEK293T cell lysates, where we identified intra- and inter-protein cross-links involving 295 proteins listed in the DisProt database [32] (Table S1). Specifically, 102 of them exhibit disordered content ranging from 20% to 100% (Table S2) including i) transcription factors/repressors and their modulators (NF-κB, YY1, TP53BP1, CREBBP), ii) histones and their associated proteins (H4C1, H3-3A, H2BC11, HDAC1, EP300), iii) proteins involved in RNA binding/processing and splicing factors (SNRPA-B, SRRM1-2, SRSF1, U2AF1), and iv) ribosomal proteins/factors (SERBP1, RPLP2, RPL4, RPL24, ElF4B, ElF4EBP1). Among those, we delved into the interactome of serpine 1 mRNA-binding protein 1 (SERBP1, also known as plasminogen activator inhibitor 1 RNA-binding protein). SERBP1 is a ∼45 kDa RNA-binding protein involved in mRNA maturation, translational regulation, and various biological functions [33, 34]. SERBP1-ribosome binding promotes ribosome hibernation [35]. SERBP1 is an IDP (see Figure 6A, according to IUPRED and DISPRED disorder predictions [36, 37]) with a short α-helix comprising residues 290-300. This α-helix has been confirmed by NMR [33] and predicted by AlphaFold2 [38] with high confidence (Figure 6B). As the conformational flexibility of IDPs does not hamper their chemical reactivity, we were able to identify 10 unique SERBP1 cross-links, recapitulating the interaction of all its structural regions with 7 binding proteins (Figure 6C). Initially, we mapped the cross-link between Lys 299 of SERBP1 and Lys 116 of the 40S Ribosomal Protein S12 (RPS12). With a Cα-Cα distance of 10 Å, this cross-link aligns well with the cryo-EM structure of the 80S ribosome [39], confirming the localization of SERBP1’s sole visualized α-helix in this protein complex (pdb id 6z6m, Figure 6D). Furthermore, we found i) the N-terminal region of SERBP1 interacting with RPS3A, RPS9, RPS14 and RPS28; and ii) the C-terminal domain interacting with RPS15 and RACK1, a scaffold protein involved in SERBP1 recruitment [40]. These PPIs possess a high STRING score and some of them have been identified in previous studies [35, 41]. We observed that SERBP1 can occupy a broad surface of the 40S ribosomal subunit, sampling a large conformational ensemble at the mRNA entry channel, as well as the A and P sites of the ribosome, thereby inhibiting translation (Figure 6E). This plasticity was also observed for SERBP1 homologues [39, 42, 43] suggesting that intrinsic disorder plays a pivotal role for the regulatory functions of these hibernation factors.

**Figure 6:**
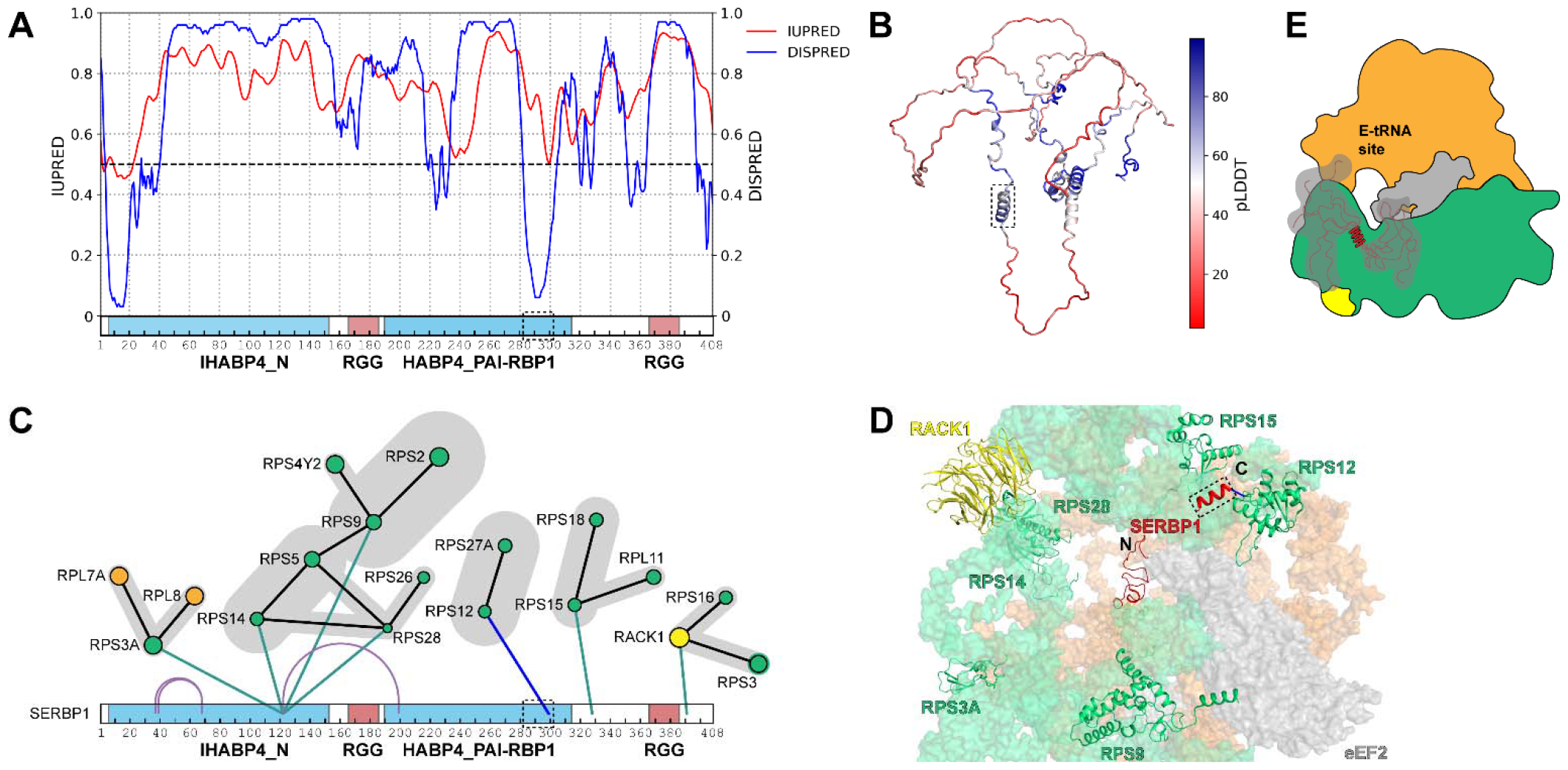
Evaluation of protein-protein interactions involving the intrinsically disordered protein SERBP1. A) IUPRED and DISOPRED disorder predictions of SERBP1, confirming its high disordered nature. The horizontal bar is displaying the SERBP1 sequence number and domain annotation. B) AlphaFold2 model of SERBP1, with the experimentally confirmed α-helix (dashed rectangle) spanning residues 290-300. C) SERBP1 interprotein cross-links to several ribosomal proteins of the 40S subunit. The cross-link involving SERBP1 solved helix (dashed rectangle) with RPS12 is colored in blue. D) Bottom view of the cryo-EM structure of the 80S ribosome (pdb id 6z6m), showing the only mappable cross-link between SERBP1 helix and RPS12. All other cross-links involved unsolved regions in either SERBP1 or ribosomal proteins. Proteins of the 40S subunit are depicted in green, proteins of the 60S subunit in orange, RACK1 in yellow, eEF2 and other associated factors in gray. RNA is not shown for clarity. E) Schematic cartoon representation of the dynamic nature of the interaction between SERBP1 and ribosome. The high disorder content allows SERBP1 to sample a vast ensemble of conformations; SERBP1 binds to the mRNA entry channel and the A and P sites of the small 40S subunit (where it interacts with eEF2, in gray), inhibiting translation.

## Conclusions

We synthesized and applied DSSI as the first imide-based cross-linker for protein 3D-structural analysis and proteome-wide interaction studies. This novel class of cross-linkers harbours an affinity tag for affinity purification of cross-links. In particular, DSSI maintains the NHS (N-hydroxysuccinimide) ester warheads and the spacer length of the widely used DSBU, but possesses a phosphonate handle for IMAC or TiO_2_ cross-link enrichment. We deciphered the gas-phase chemistry of DSSI and parametrized its MS-cleavability for automated cross-link identification. Its diagnostic fragment ion doublets can be recognized by the MeroX software for a fast and reliable identification of cross-links in highly complex samples. Furthermore, we demonstrated the reactivity of DSSI by cross-linking α-syn and HEK293T cell lysates. In total, we identified 5,602 unique cross-links involving 1,241 proteins. DSSI targets proteins in a wide range of cellular compartments and expands the scope of XL-MS. DSSI cross-links were validated based on the high-resolution structures of three large protein complexes, the CCT complex, the 26S proteasome, and the 80S ribosome. The Cα-Cα distance distribution of DSSI cross-links matches that of DSBU. Beyond characterizing PPIs in HEK293T cells, we derived morphological insights into IDPs, such as SERBP1. We anticipate that the MS-cleavable DSSI and other imide-based cross-linkers, in combination with SEC and IMAC enrichment of cross-link products, will contribute to unveiling structural details of the cell interactome. Future studies will be directed to address the IMAC enrichment.

## ASSOCIATED CONTENT

### Supporting Information

The Supporting Information is available free of charge on the ACS Publications website.

Experimental details (Synthesis and characterization of DSSI, cross-linking, enzymatic digestion, SEC, and data analysis) (PDF). Tables of cross-linked IDPs (XLSX). MS data have been deposited to the ProteomeXchange Consortium via the PRIDE partner reposi-tory with the project accession PXD050960, username; password: Id1BhOxl

## AUTHOR INFORMATION

### Authors

**Alessio Di Ianni** - Department of Pharmaceutical Chemistry and Bioanalytics, Institute of Pharmacy, Martin Luther University Halle-Wittenberg, Kurt-Mothes-Str. 3, D-01620 Halle/Saale, Germany, Center for Structural Mass Spectrometry, Martin Luther University Halle-Wittenberg, Kurt-Mothes-Str. 3, D-01620 Halle/Saale, Germany

**Christian H. Ihling** - Department of Pharmaceutical Chemistry and Bioanalytics, Institute of Pharmacy, Martin Luther University Halle-Wittenberg, Kurt-Mothes-Str. 3, D-01620 Halle/Saale, Germany, Center for Structural Mass Spectrometry, Martin Lu-ther University Halle-Wittenberg, Kurt-Mothes-Str. 3, D-01620 Halle/Saale, Germany

**Andrea Sinz** - Department of Pharmaceutical Chemistry and Bioanalytics, Institute of Pharmacy, Martin Luther University Halle-Wittenberg, Kurt-Mothes-Str. 3, D-01620 Halle/Saale, Germany, Center for Structural Mass Spectrometry, Martin Luther University Halle-Wittenberg, Kurt-Mothes-Str. 3, D-01620 Halle/Saale, Germany

**Tomáš Vranka** – CF Plus Chemicals s.r.o., Karásek 1767/1, 621 00 Brno-Řečkovice, Czechia

**Václav Matoušek** - CF Plus Chemicals s.r.o., Karásek 1767/1, 621 00 Brno-Řečkovice, Czechia

### Author Contributions

The manuscript was written through contributions of all authors. / All authors have given approval to the final version of the manuscript.

## ACKNOWLEDGMENT

CI acknowledges financial support by the Italian Ministry of University and Research (MUR) (PRIN 2022 – Project 20225HNCZK) and the European Union — Next Generation EU (PRIN 2022 PNRR – Project P20224WAME). AS acknowledges financial support by the DFG (RTG 2467, project number 391498659 “Intrinsically Disordered Proteins-Molecular Principles, Cellular Functions, and Diseases”, CRC 1423, project number 421152132), INST 271/404-1 FUGG, INST 271/405-1 FUGG), the Federal Ministry for Economic Affairs and Energy (BMWi, ZIM project KK5096401SK0), the region of Saxony-Anhalt, and the Martin Luther University Halle-Wittenberg (Center for Structural Mass Spectrometry). The authors thank Frank Hause for their time and constructive input invested into this work. The authors are indebted to Dr. Marcel Köhn and his lab for providing HEK293T cells and assistance.

